# Are *Escherichia coli* causing recurrent cystitis just ordinary Uropathogenic *E. coli* (UPEC) strains?

**DOI:** 10.1101/2023.11.08.566351

**Authors:** Nicolas Vautrin, Sandrine Dahyot, Marie Leoz, François Caron, Maxime Grand, Audrey Feldmann, François Gravey, Stéphanie Legris, David Ribet, Kévin Alexandre, Martine Pestel-Caron

## Abstract

Specific determinants associated with Uropathogenic *Escherichia coli* (UPEC) causing recurrent cystitis are still poorly characterized. The aims of this study were (i) to describe genomic and phenotypic traits associated with recurrence using a large collection of recurrent and paired sporadic UPEC isolates, and (ii) to explore within-host genomic adaptation associated with recurrence using series of 2 to 5 sequential UPEC isolates. Whole genome comparative analyses between 24 recurrent cystitis isolates (RCIs) and 24 phylogenetically paired sporadic cystitis isolates (SCIs) suggested a lower prevalence of putative mobile genetic elements (MGE) in RCIs, such as plasmids and prophages. The intra-patient evolution of the 24 RCI series over time was characterized by SNP occurrence in genes involved in metabolism or membrane transport, and by plasmid loss in 5 out of the 24 RCI series. Genomic evolution occurred early in the course of recurrence, suggesting rapid adaptation to strong selection pressure in the urinary tract.

However, RCIs did not exhibit specific virulence factor determinants and could not be distinguished from SCIs by their fitness, biofilm formation, or ability to invade HTB-9 bladder epithelial cells. Taken together, these results suggest a rapid but not convergent adaptation of RCIs that involves both strain- and host-specific characteristics.

**Author summary:** The recurrence of cystitis is a frequent but poorly understood phenomenon. There are currently many hypotheses trying to explain recurrence, but data on large collections of well-characterized clinical isolates are lacking. In order to identify specific recurrence-associated markers, we conducted a large genomic and phenotypic study involving 48 well-characterized cystitis isolates: 24 recurrent cystitis isolates (RCIs) and 24 pairs of isolates causing sporadic cystitis (SCIs). Moreover, we were able to explore intra-host overtime RCI evolution, by analyzing up to 5 sequential UPEC isolates per RCI series. Our results suggest that RCI rapidly adapt to their host through mobile genetic elements loss and SNP accumulation in genes involved in metabolism and membrane transport. However, no convergent genomic nor phenotypic evolution was observed between isolates collected from distinct patients. Taken together, these results suggest a host-shaped evolution of RCIs, highlighting a need for future studies focused on the host-pathogen relationships.

## Introduction

Urinary tract infections (UTIs) are very common bacterial infections in women as more than half of them will develop at least one UTI during their lifetime [1]. Up to 25% of women who had a UTI will experience a second one within a year [2]. If a woman presents more than 2 episodes within a 6-month period or 3 episodes within 12 months, she will be considered as suffering from recurrent UTI (rUTI) [3]. rUTI is a public health concern, and is associated with an economic, societal, and personal burden. It represents 1% to 6% of all medical visits and is the second leading cause of antibiotic consumption in the United States [2]. Its annual cost is estimated at 1.6 billion US dollars [4]. Moreover, rUTI negatively impacts quality of life, as they promote anxiety and depression [4].

rUTIs are mainly caused by uropathogenic *Escherichia coli* (UPEC) [2] whether relapses or reinfections. Relapse is defined by infection with the same strain as the initial infecting strain whereas reinfection corresponds to infection with a strain different from the initial one [5]. Using pulsed-field gel electrophoresis (PGFE), it was previously estimated that 47 to 81% of UPEC rUTI were due to relapses [3]. However, this statement was primarily based on typing methods such as serotyping [6] and pulsed-field gel electrophoresis (PFGE) [5,7,8] which have limited discriminatory power compared to molecular-based methods [9]. Using next generation sequencing (NGS) and CH typing - a molecular typing method based on the polymorphism of internal fragments of two genes (*fumC* and *fimH*) [10], we recently observed that the frequency of relapses was only 30.6% [11].

UPEC virulence primarily relies on their ability to survive and grow in urine, as well as to adhere to and invade urothelial cells [12]. Although certain mechanisms involved in UPEC virulence have been well described, the physiopathology of recurrent cystitis remains poorly characterized [13]. Among the existing hypotheses, it has been postulated that the ability of UPECs to persist in the bladder by forming intracellular bacterial communities (IBC) and quiescent intracellular reservoir (QIR) is involved in recurrent cystitis [14–16]. However, this phenomenon has not yet been explored in large cohorts of patients. Moreover, no specific genomic nor phenotypic marker have been identified to distinguish UPECs associated with recurrent cystitis from those causing sporadic cystitis [17–19].

In this context, our study first aimed to identify genomic and/or phenotypic determinants associated with recurrence using a collection of 24 recurrent cystitis isolates (RCIs) and 24 phylogenetically paired isolates responsible for sporadic cystitis (SCIs), sampled from patients with clinically well-characterized cystitis over a 17-month period (VITALE study, NTC02292160). The second aim was to describe within-host microevolution of the 24 RCIs over time, by analysing the genomic and phenotypic changes in 24 series of 2 to 5 sequential isolates.

## Results

### Analysis of 26 virulence factors determinants did not distinguish RCIs from SCIs

To identify genomic traits associated with recurrence, we compared the genomes of 24 initial RCI (iRCIs) with those of 24 SCIs that phylogenetically matched based on CH typing (Fig 1, Table S1). The presence and protein sequence of 26 virulence factor determinants (VFDs) of interest [17] were compared between these two groups (Table 1).

**Fig 1.**
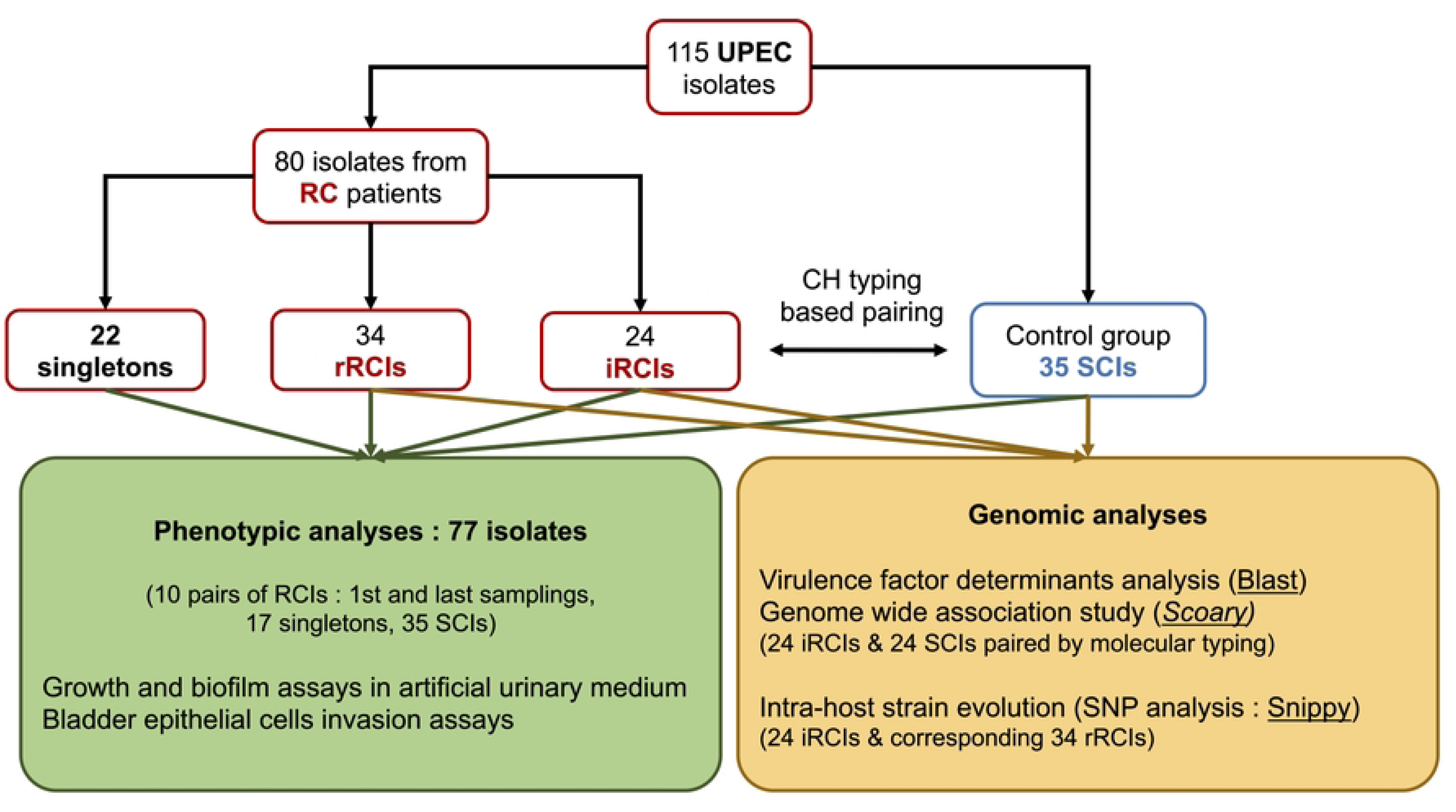
Isolate and workflow description. RC patients = patients with recurrent cystitis, iRCI = initial recurrent cystitis isolate, rRCI = recurrent cystitis isolate associated with relapse(s), SCI = sporadic cystitis isolate.

**Table 1.**
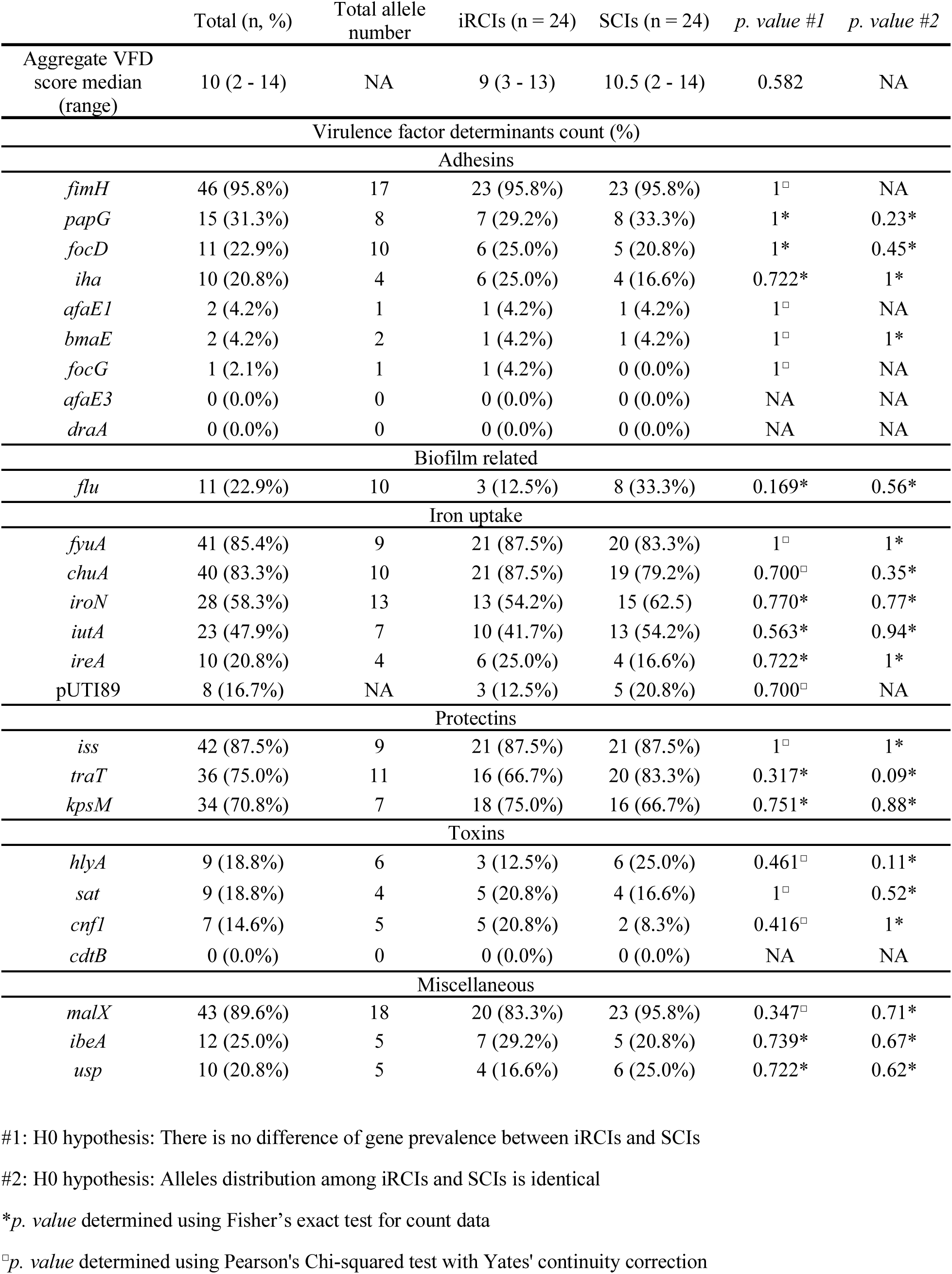
Prevalence of virulence factors determinants (VFDs) in the 24 iRCIs and 24 SCIs. NA stands for “non-applicable”.

The prevalence of the 26 VFDs ranged from 0.0% to 95.8% among the 48 UPEC isolates, with no significant difference between the iRCI and SCI groups. The aggregate VFD score median, determined as previously described by Ejrnaes *et al.* [17], was also not significantly different between the two groups of isolates (median of 9 for iRCIs *vs.* 10 for SCIs, *p* = 0.665) (Table 1). Moreover, a principal component analysis based on the detection of the 26 VFDs revealed that iRCIs and SCIs did not cluster separately (Fig 2). There were up to 18 alleles identified per gene (*malX*), with no significant association with recurrence (Table 1, Fig S1).

**Fig 2.**
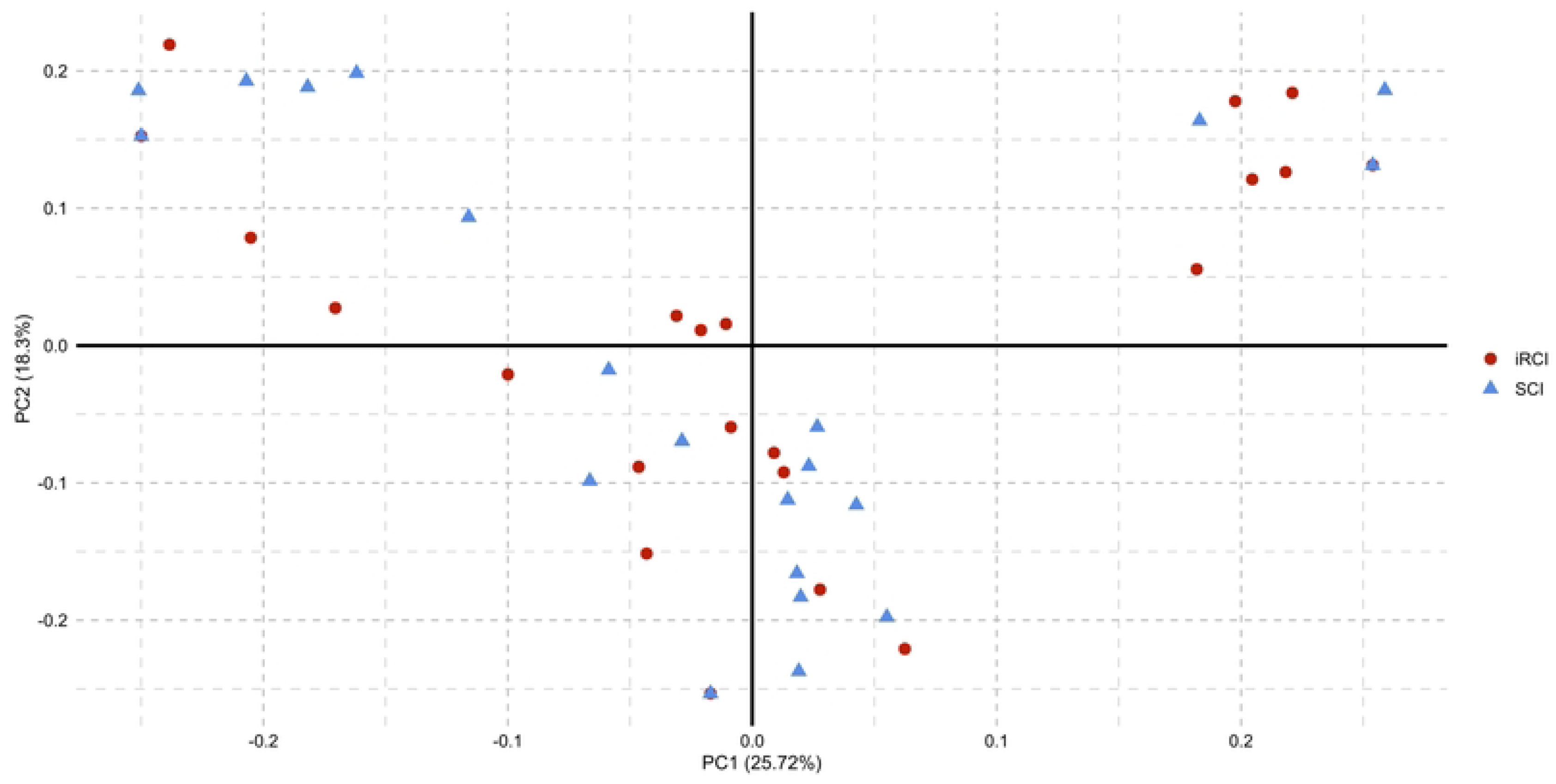
Principal component analysis based on the detection of virulence factor genes. Red dots represent iRCIs (initial recurrent cystitis isolates) and blue triangles represent SCIs (sporadic cystitis isolates).

Among the 10 adhesin-encoding genes investigated, *fimH* was the most prevalent (95.8%) (Table 1). We did not test the association of the 17 *fimH* alleles with clinical context since this gene belongs to the typing scheme used to pair SCIs to the 24 iRCIs. The second most common gene in this virulence group was *papG* (31.3%). Two of the four previously described *papG* variants [20,21] were identified (*papGII* and *papGIII*), although they were not significantly associated with the RCI or SCI groups.

The most prevalent markers associated with iron uptake were *fyuA* (85.4%) and *chuA* (83.3%), while the least frequent was pUTI89, a plasmid associated with iron uptake [22]. The great majority of the isolates (41/48, 85.4%) presented at least two iron uptake genes. Three isolates (one iRCI/SCI pair and one SCI) were negative for all iron uptake markers tested.

The global prevalence of toxin-encoding genes (*hlyA*, *sat*, *cnf1, cdtB*) and of the *flu* gene associated with biofilm production was low (<25%), while the prevalence of genes encoding protectins was high (>70%).

### iRCI genomes were significantly smaller than phylogenetically paired SCIs

The mean genome size of iRCIs was significantly lower than that of SCIs (5.01 Mb *vs*. 5.11 Mb, respectively, *p* < 0.05), resulting in a significantly lower mean protein count (4,658 *vs*. 4,776 in iRCIs *vs*. SCIs, respectively, *p* < 0.05). The pan-genome of the 48 assemblies contained 15,682 genes, of which 2,864 (18.3%) were considered core genes. Phylogenetic analysis of the core genes confirmed that isolates clustered based on CH type rather than on the sporadic/recurrent clinical context (Fig 3A), consistent with the SCI/iRCI CH type-based pairing strategy.

**Fig 3.**
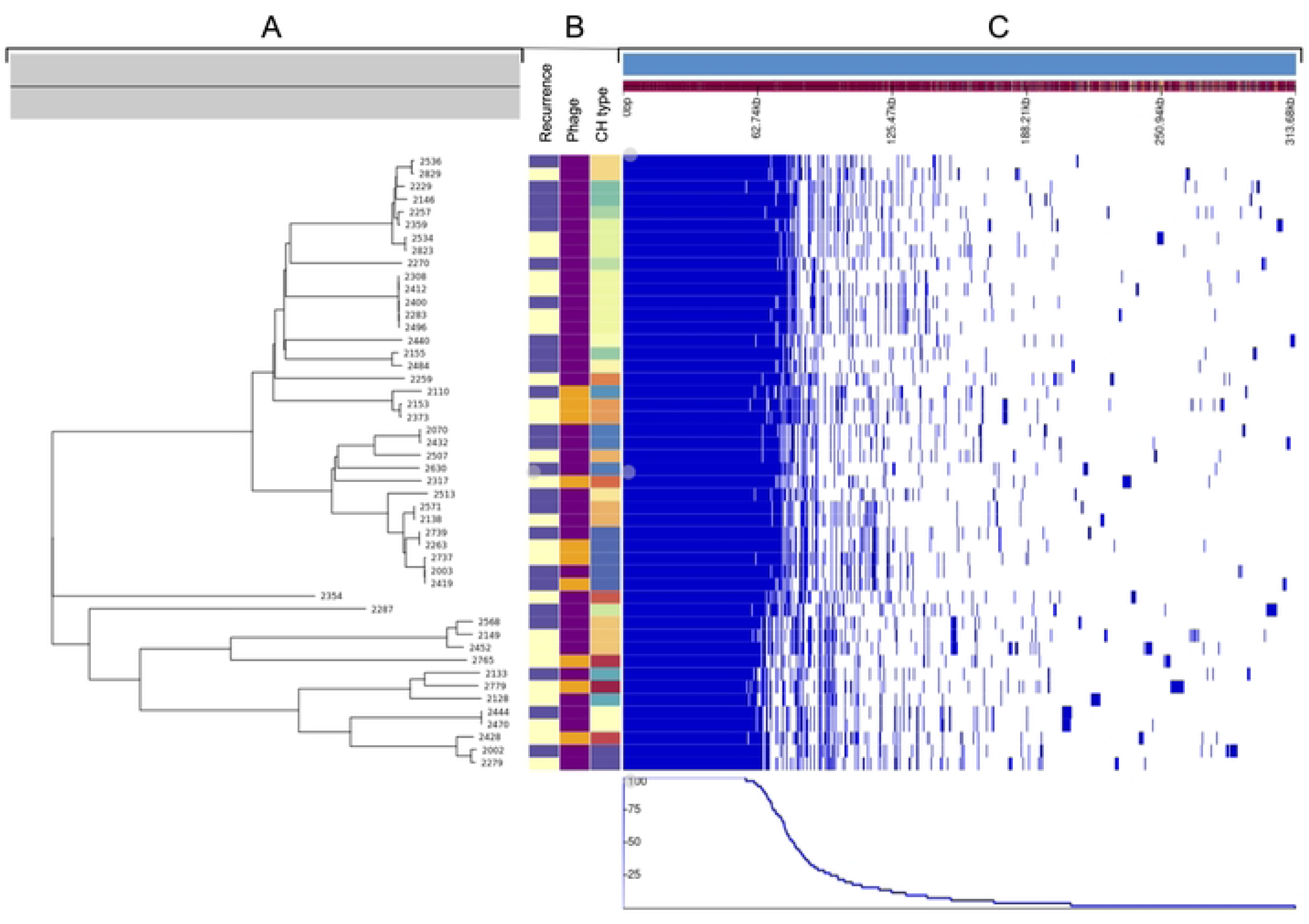
Core genome analysis of the 24 iRCIs and their 24 CH type paired SCIs. iRCIs = initial recurrent cystitis isolates, SCIs = sporadic cystitis isolates. **A.** Neighbor-Joining phylogenetic tree based on the 48 strains’ translated core gene alignment (2,755,382 amino acid positions). **B.** Color-coded CH type, prophage presence (purple = absent, yellow = present), clinical context of recurrence (purple = iRCI, yellow = SCI) for the 48 strains. **C.** Presence/absence (blue/white) of the 15,682 genes from the pan genome in each strain, sorted by gene frequency among the 48 strains. The first 2,864 genes on the left represent the core genome.

A genome-wide association study (GWAS) was performed based on the presence or absence of accessory genes in iRCIs *vs.* SCIs (Fig 3C). Only two genes were significantly less frequent in RCI genomes (Sgene1, *p* = 0.009 and Sgene2, *p* = 0.009). Proteins encoded by these genes were annotated as phage-associated proteins by InterProScan. Sgene1 encoded an unknown DNA binding protein with a helix turn helix domain (LocusTag OMANCKEL_0742 in CS2737) and Sgene 2 encoded a regulatory phage protein from the Cox family (LocusTag OMANCKEL_0743 in CS2737; InterPro entry: IPR019679). In the 7 SCIs where Sgene1 and Sgene2 were identified, they were located back-to-back and followed by a sequence of up to 40 hypothetical proteins-coding genes, most of which being annotated as phage-associated proteins by InterProScan. This putative prophage was located between the *cpx* operon and the *fieF* gene (also named *yiiP*), encoding an envelope stress response system and a ferrous iron efflux pump, respectively.

Systematic Phigaro analysis of the 48 draft genomes confirmed the presence of diverse phage sequences inserted between the *cpx* operon and *fieF* gene of 8 SCIs, but in only 2 of their paired iRCIs (Fig 3B).

iRCI reads were mapped to their paired SCI assembly to illustrate the phage presence/absence (Fig 4A). This analysis also highlighted that some very large SCI contigs (up to 40 kb) were absent in paired iRCI; some of these contigs contained genes from the *tra* operon, which suggested that they derived from plasmids (Fig 4B).

**Fig 4.**
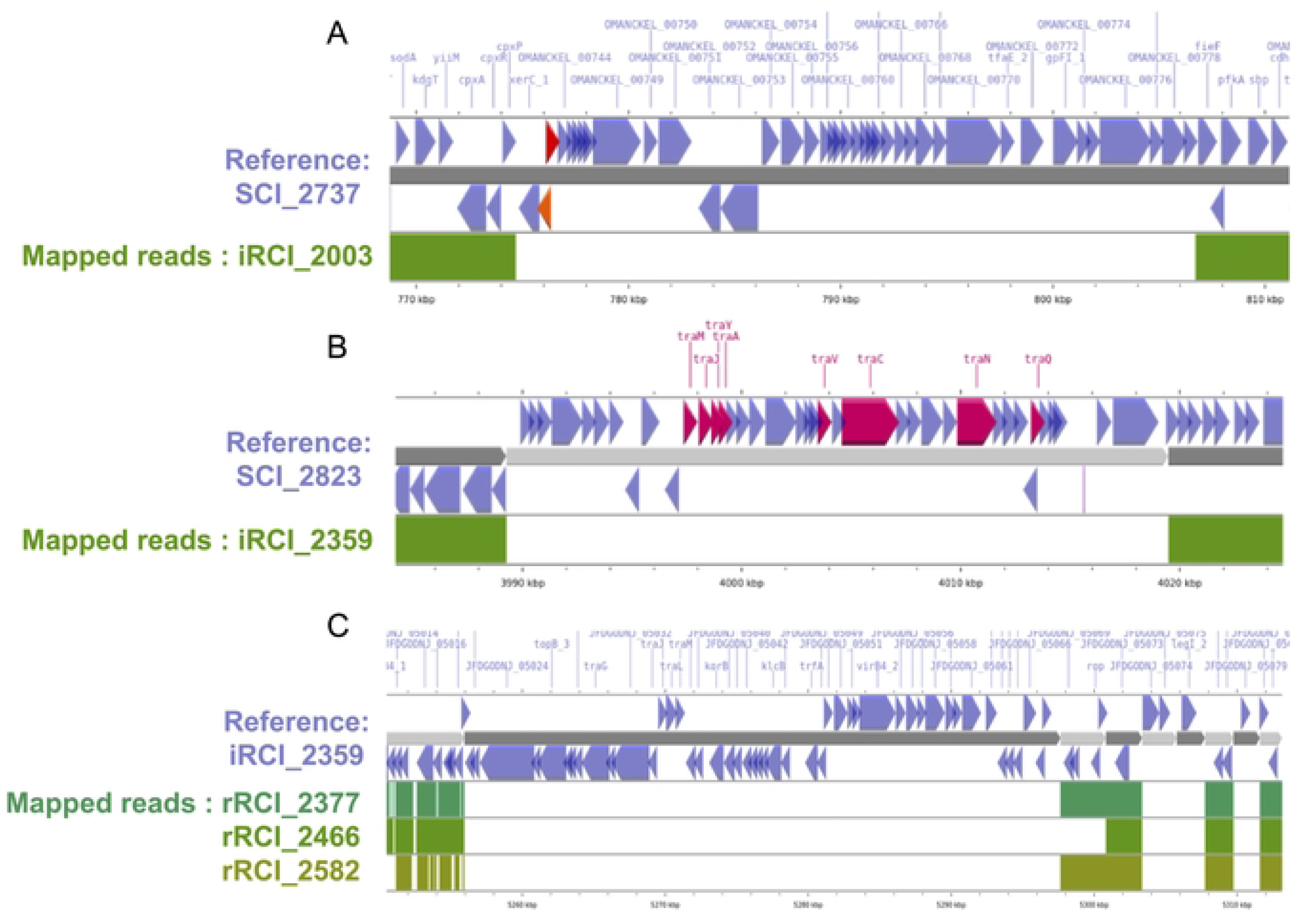
Examples of mobile genetic elements that are found in SCIs but not in their paired iRCIs (A and B) or lost in sequential relapses in RCI patients (C). Contigs from genome assemblies are shown in an alternance of dark and light grey arrows, iRCIs = initial recurrent cystitis isolates, rRCI = recurrent cystitis isolate associated with relapse(s), SCIs = sporadic cystitis isolates. Just above and below, blue arrows indicate the location and orientation of CDSs on the contigs. **A.** A 40 kb portion of SCI_2737 contig #6 is represented in dark grey. Orange and red arrows respectively indicate SCI-associated genes Sgene1 and Sgene2. Beneath, the regions covered by the reads from paired iRCI_2003 are indicated in green. The >30 kb SCI_2737 genome region that is not covered by iRCI_2003 reads includes 44 phage proteins similar to those from Peduovirus P24B2 (AccNum NC_049387). **B.** SCI_2823 contig #59 is represented in light grey. Pink arrows indicate the location of plasmid-specific genes from the *tra* operon. The reads from paired iRCI_2359 map to SCI_2823 contigs #58 and #60, but not to the >30 kb contig #59, which derives from a plasmid similar to the *Escherichia coli* strain HS13-1 plasmid pHS13-1-IncF (AccNum CP026494) **C.** Contig #3 from hybrid genome assembly of iRCI_ 2359 is indicated in dark grey. Below, the coverage of this contig by the reads of the corresponding rRCIs is shown in shades of green. iRCI_2359 contig #3 corresponds to a 42kb plasmid similar to the *E. coli* pRHB15-C18_3, (AccMum CP057780.1) that is lost in the first to third relapses.

Taken together, these results suggested that mobile genetic elements (MGEs) were less frequent in iRCIs than in SCIs, consistent with their lower genome sizes.

### Longitudinally sampled RCIs lost plasmids over the course of relapses

Long-read whole genome sequencing and hybrid assemblies were performed on the 24 iRCIs investigated so far in order to better describe their genomes and in particular their plasmids. Hybrid assembly reduced by 19-fold the mean number of contigs per genome and modestly increased the total genome size (mean of 5.10 Mb instead of 5.01 Mb), though mean estimated completeness remained 99.9% for hybrid as well as short read assemblies according to BUSCO.

To longitudinally investigate RCI evolution over time, short reads from each recurrent cystitis isolate associated with relapse(s) (rRCI, one to four relapses per patient) were mapped to the hybrid assembly of their iRCI, used as an intra-host reference. Interestingly, 5 of the 24 RCIs (21%) lost up to 3 circular contigs over the course of relapses, identified as plasmids using blast. Of the 8 lost plasmids, 4 were small cryptical plasmids (<5kb) coding only hypothetical proteins (Table S2). The three others were large plasmids (>40kb) encoding conjugation system (*tra* operon), toxin-antitoxins systems, and a few proteins that could not be associated with urovirulence. In 7 cases out of 8, plasmid loss occurred between the iRCI and the first rRCI (exemplified in Fig 4C).

### Non-synonymous SNPs of longitudinally sampled RCIs often occurred in genes involved in metabolism and membrane transport

To investigate RCI micro-evolution, we looked for SNP acquisition over time. In the 24 relapse series, a total of 666 SNPs were identified. Among these, 255 (38.3%) were non-synonymous SNPs (nsSNPs). Functional annotation of the 160 proteins affected by these nsSNPs was successful for 107 proteins, 58 of which were mapped in defined biological processes by blastKoala (Table 2). These 58 proteins were mostly involved in diverse metabolic pathways (*n* = 22, 37.9%) and membrane transport (*n* = 12, 20.7%).

**Table 2.** Functional classification of the 58 proteins encoded by genes in which non-synonymous SNPs have occurred overtime in the RCI series. Annotation and corresponding biological process were provided by BlastKoala. Of note, a single gene can be involved in several biological processes.

| Functionnal category | Number of genes |
| --- | --- |
| Metabolism | (*n* = 22 genes) |
| **Global and overview maps** | **17** |
| Diverses metabolic pathways | 16 |
| Biosynthesis of secondary metabolites | 9 |
| Microbial metabolism in diverse environnements | 4 |
| Carbon metabolism | 2 |
| Biosynthesis of amino acids | 1 |
| Biosynthesis of nucleotide sugars | 1 |
| Biosynthesis of cofactors | 3 |
| Biosynthesis of aromatic compounds | 1 |
| **Carbohydrate metabolism** | **5** |
| Glycolysis / Gluconeogenesis | 1 |
| Citrate cycle (TCA cycle) | 1 |
| Pentose and glucuronate interconversions | 1 |
| Fructose and mannose metabolism | 1 |
| Amino sugar and nucleotide sugar metabolism | 1 |
| Pyruvate metabolism | 2 |
| Propanoate metabolism | 2 |
| Butanoate metabolism | 1 |
| C5-Branched dibasic acid metabolism | 1 |
| **Energy metabolism** | **2** |
| Carbon fixation pathways | 2 |
| Methane metabolism | 1 |
| **Lipid metabolism** | **1** |
| Fatty acid degradation | 1 |
| **Nucleotide metabolism** | **1** |
| Purine metabolism | 1 |
| **Amino acids metabolism** | **5** |
| Glycine, serine and threonine metabolism | 1 |
| Lysine degradation | 1 |
| Arginine and proline metabolism | 1 |
| Histidine metabolism | 1 |
| Tyrosine metabolism | 1 |
| **Metabolism of other amino acids** | **2** |
| Taurine and hypotaurine metabolism | 1 |
| Gluthathione metabolism | 1 |
| **Glycan biosynthesis and metabolism** | **5** |
| Lipopolysaccharide biosynthesis | 4 |
| Exopolysaccharide biosynthesis | 1 |
| **Metabolism of cofactors and vitamins** | **3** |
| Biotin metabolism | 1 |
| Porphyrin metabolism | 1 |
| Ubiquinone and other terpenoid-quinones biosynthesis | 1 |
| **Metabolism of terpenoids and polyketides** | **1** |
| Biosynthesis of siderophore group nonribosomal peptides | 1 |
| **Biosynthesis of other secondary metabolites** | **1** |
| Tropane, piperidine and pyridine alkaloid biosynthesis | 1 |
| **Xenobiotics biodegradation and metabolism** | **4** |
| Chloroalkane and chloroalkene degradation | 1 |
| Naphthalene degradation | 1 |
| Metabolism of xenobiotics by cytochrome P450 | 1 |
| Drug metabolism - cytochrome P450 | 1 |
| Drug metabolism - other enzymes | 1 |
| **Environnemental information processing** | **(n = 15 genes)** |
| **Membrane transport** | **11** |
| ABC transporters | 10 |
| Bacterial secretion systems | 1 |
| **Signal transduction** | **3** |
| Two-component system | 3 |
| **Genetic information processing** | **(n = 6 genes)** |
| **Translation** | **2** |
| Ribosome | 1 |
| Aminoacyl-tRNA biosynthesis | 1 |
| **Folding, sorting and degradation** | **3** |
| RNA degradation | 3 |
| **Replication and repair** | **4** |
| DNA replication | 2 |
| Base excision repair | 1 |
| Mismatch repair | 2 |
| Homologous recombination | 3 |
| **Cellular processes** | **(n = 6 genes)** |
| **Cellular community – prokaryotes** | **5** |
| Quorum sensing | 2 |
| Biofilm formation – *Vibrio cholerae* | 2 |
| Biofilm formation – *Escherichia coli* | 2 |
| **Cell motility** | **1** |
| Flagellar assembly | 1 |
| **Human diseases** | **(n = 5 genes)** |
| **Cancer: overview** | **1** |
| Pathways in cancer | 1 |
| Chemical carcinogenesis – DNA adducts | 1 |
| Chemical carcinogenesis – receptor activation | 1 |
| Chemical carcinogenesis – reactive oxygen species | 1 |
| **Cancer: specific types** | **1** |
| Hepatocellular carcinoma | 1 |
| **Infectious disease: bacterial** | **1** |
| Shigellosis | 1 |
| Yersinia infection | 1 |
| Pertussis | 2 |
| Bacterial invasion of epithelial cells | 1 |
| **Infectious disease: parasitic** | **1** |
| Amoebiasis | 1 |
| **Cardiovascular disease** | **1** |
| Fluid shear and atherosclerosis | 1 |
| **Drug resistance: antineoplastic** | **1** |
| Platinum drug resistance | 1 |
| **Organismal systems** | **(*n* = 1 gene)** |
| **Aging** | **1** |
| Longevity regulating pathway – worm | 1 |

Notably, 10 out of the 24 RCIs acquired nsSNPs in genes encoding diverse ABC membrane transporters. Among these, two were involved in metal ion transport (nickel import ATP-binding protein NikD, LocusTag = OKMJCHAP_00349 in iRCI_2002; and Fe(3+) dicitrate transport system permease protein FecC, LocusTag = AHEJCLJF_03297 in iRCI_2110). One membrane transport protein (lipopolysaccharide export system ATP-binding protein LptB) acquired nsSNPs in two distinct relapse series (LocusTag = MMGADFLO_03138 in iRCI_2229 and HFPIPPMJ_01315 in iRCI_2359). Of note, another nsSNP occurred in a gene involved in lipopolysaccharide biosynthesis (lipid A export ATP-binding/permease protein MsbA, LocusTag = AKCFODAC_00517 in iRCI_2484).

Overtime conserved SNPs were investigated in the 7 RCI series that contained more than 2 episodes. A total of 13 SNPs identified in the first rRCI were conserved in the following ones. Ten of these overtime conserved SNPs occurred in ORFs and 6 of these were non synonymous. Of note, two genes that acquired conserved nsSNPs were involved in membrane transport according to Prokka: an ABC transporter (Inner membrane ABC transporter permease protein YdcV, LocusTag = KNBHMFIB_00972 in iRCI_2287) and a permease involved into peptide transport (Dipeptide and tripeptide permease A, LocusTag = MMGADFLO_01450 in iRCI_2229). Other genes affected by overtime-conserved nsSNPs encoded ribonucleases and a sigma-E factor regulatory protein (Table S3).

### RCI genomic evolution rate tended to decrease overtime

Evolution rates between each rRCI and its corresponding iRCI ranged from 0 to 1.67 SNPs/day (mean of 0.27 SNP/day). Evolution rates tended to decrease over time, even if this trend was not significant (*p* = 0.124) (Fig 5A). However, an outlier (circled in red on Fig 5A) acquired 378 SNPs over a long period (226 days), 93% of which SNPs were located in only two plasmid-derived contigs. In depth comparison of the iRCI and rRCI contigs from this outlier suggested that these SNPs were artifacts due to the comparison of similar regions that derived from different plasmids. When discarding this outlier, the evolution rate decrease over time became significant (*p* = 0.028).

**Fig 5.**
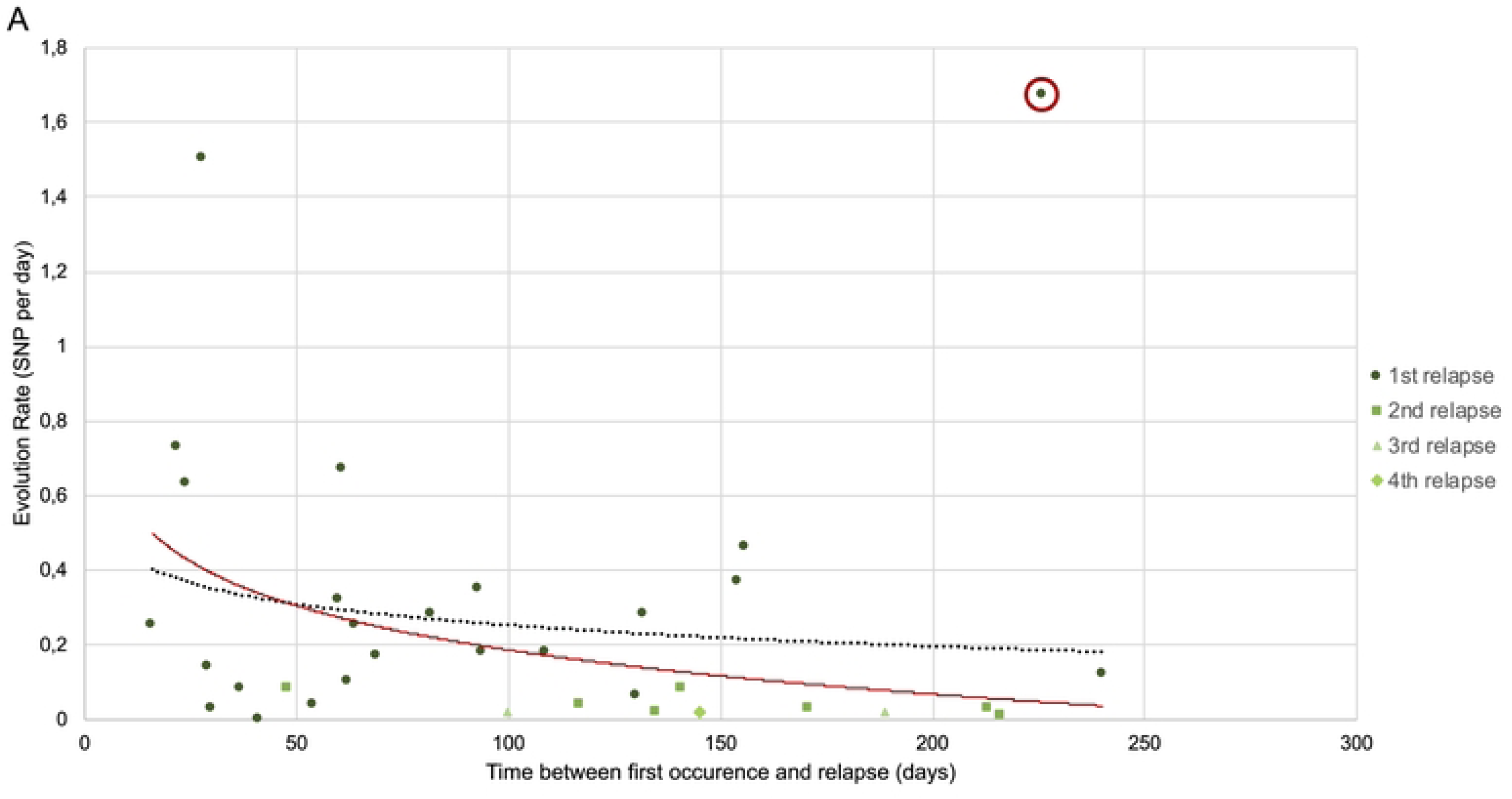
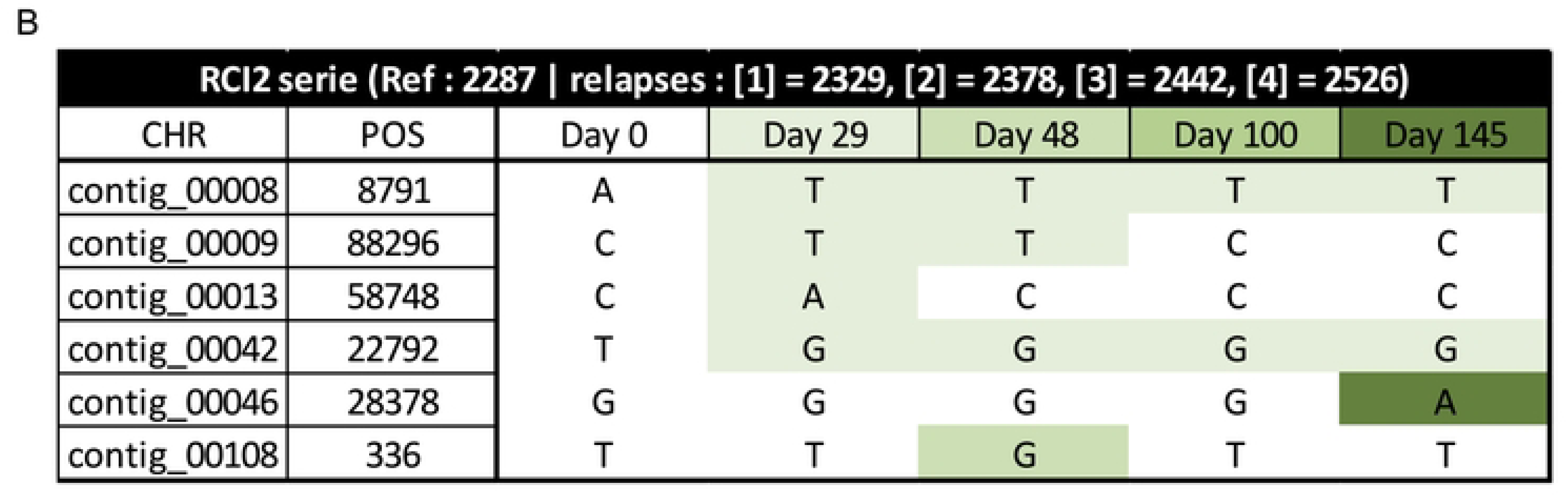
Intra patient micro-evolution of RCIs by SNP acquisition. RCIs **=** recurrent cystitis isolates **A.** Global representation of the evolution rates (SNPs per day) from the 24 series of intra-patient RCIs depending on the time elapsed between the first occurrence and the relapse (days). An outlier is circled in red. The black dotted curve represents the non-significant trend when including all the dots (*p* = 0.124). The red curve represents the significant trend when excluding the outlier (*p* = 0.028). **B.** Individual representation of the core SNPs observed over time in the largest intra-patient RCI serie (RCI2). SNPs are color-coded depending of the time of emergence (the darker, the latter). CHR = Contig on which the SNP was identified, POS = position of the SNP on the contig.

This trend was also observed at the individual scale within the largest RCI serie (RCI2), with four relapses): most of the overtime-conserved SNPs occurred early, often between the iRCI and the first rRCI. Afterwards, a dynamic of SNP emergence and clearance was observed, as shown in Fig 5B.

### Growth in AUM did not discriminate RCIs from SCIs

The growth of 72 isolates was evaluated in rich medium (lysogenic broth [LB]) and in artificial urinary medium (AUM). These isolates included 10 RCI pairs (10 iRCIs and their corresponding 10 last rRCIs), 35 SCIs and 17 singletons isolates (isolates responsible for a single infection in patients with recurrent cystitis) (Fig 1). The mean doubling times of the four groups (iRCI, rRCI, SCI and singletons) were similar in LB (G_iRCI_ = 22.9 ± 1.25min, G_rRCI_ = 23.2 ± 1.79min, G_SCI_ = 23.0 ± 1.49min and G_Singleton_ = 23.6 ± 1.99min; *p*=0.215) and not statistically different from those of the reference strains UTI89 and K12 (G_UTI89_ = 23.7 ± 1.25min and G_K12_ = 23.2 ± 0.97min) (Fig 6A). In AUM, all isolates grew significantly slower (*p* < 0.01) than in LB (G_iRCI_ = 43.6 ± 3.50min, G_rRCI_ = 43.6 ± 2.18min, G_SCI_ = 46.2 ± 4.24min and G_Singleton_ = 47.9 ± 6.34min) (Fig. 6B). Even if intra-group variability was higher in AUM than in LB, no significant differences were observed between groups and *E. coli* UTI89 in AUM (G_UTI89-AUM_ = 43.3 ± 2.53min). However, the laboratory strain *E. coli* K12, grew significantly slower than the other isolates (G_K12-AUM_ = 96.6 ± 11.6min, *p* < 0.01) (Fig 6B). Furthermore, we observed only one significant growth rate increase in RCI2 pairs that could not be link to a particular genomic evolution (Fig S2).

**Fig 6.**
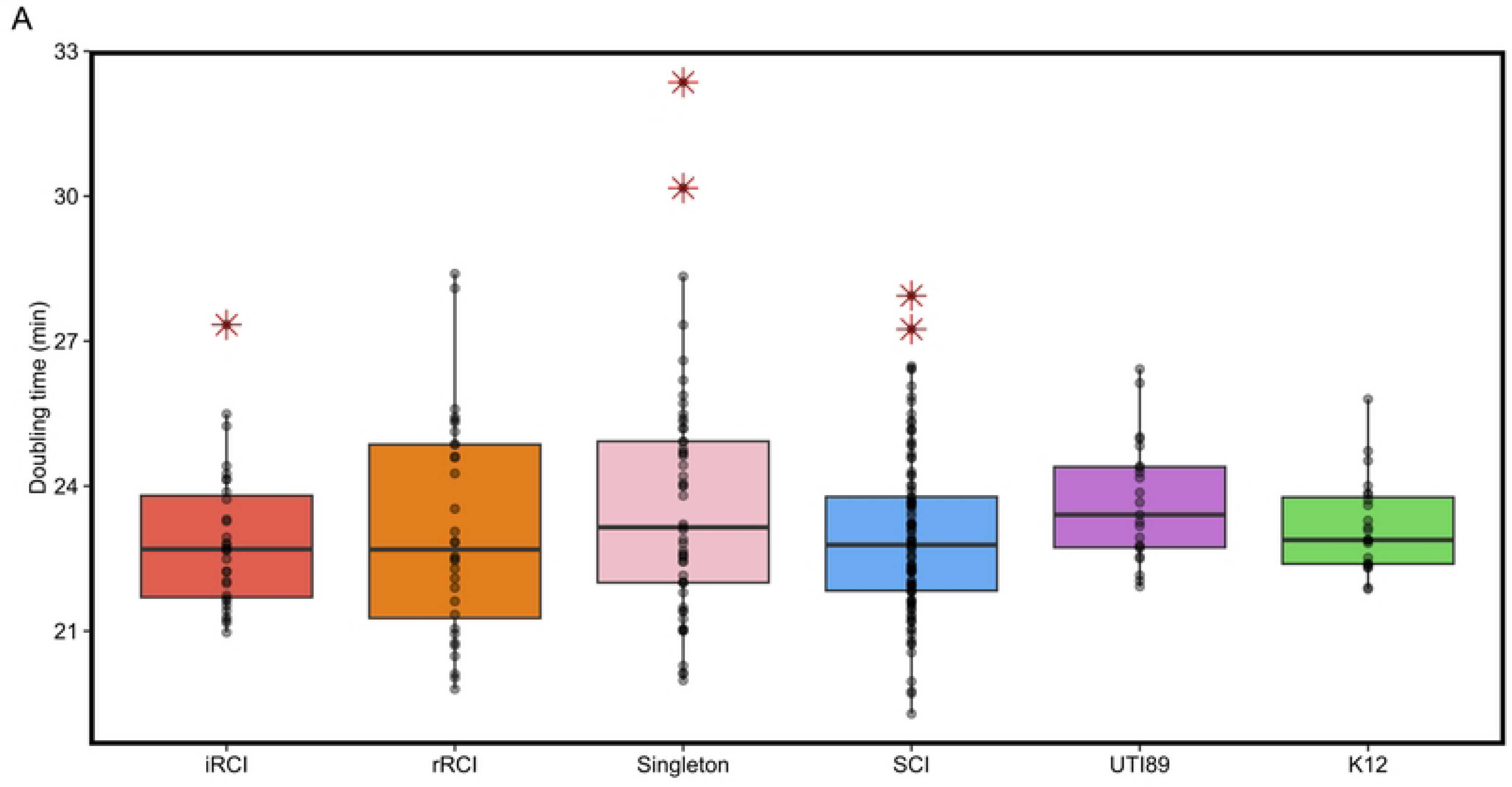
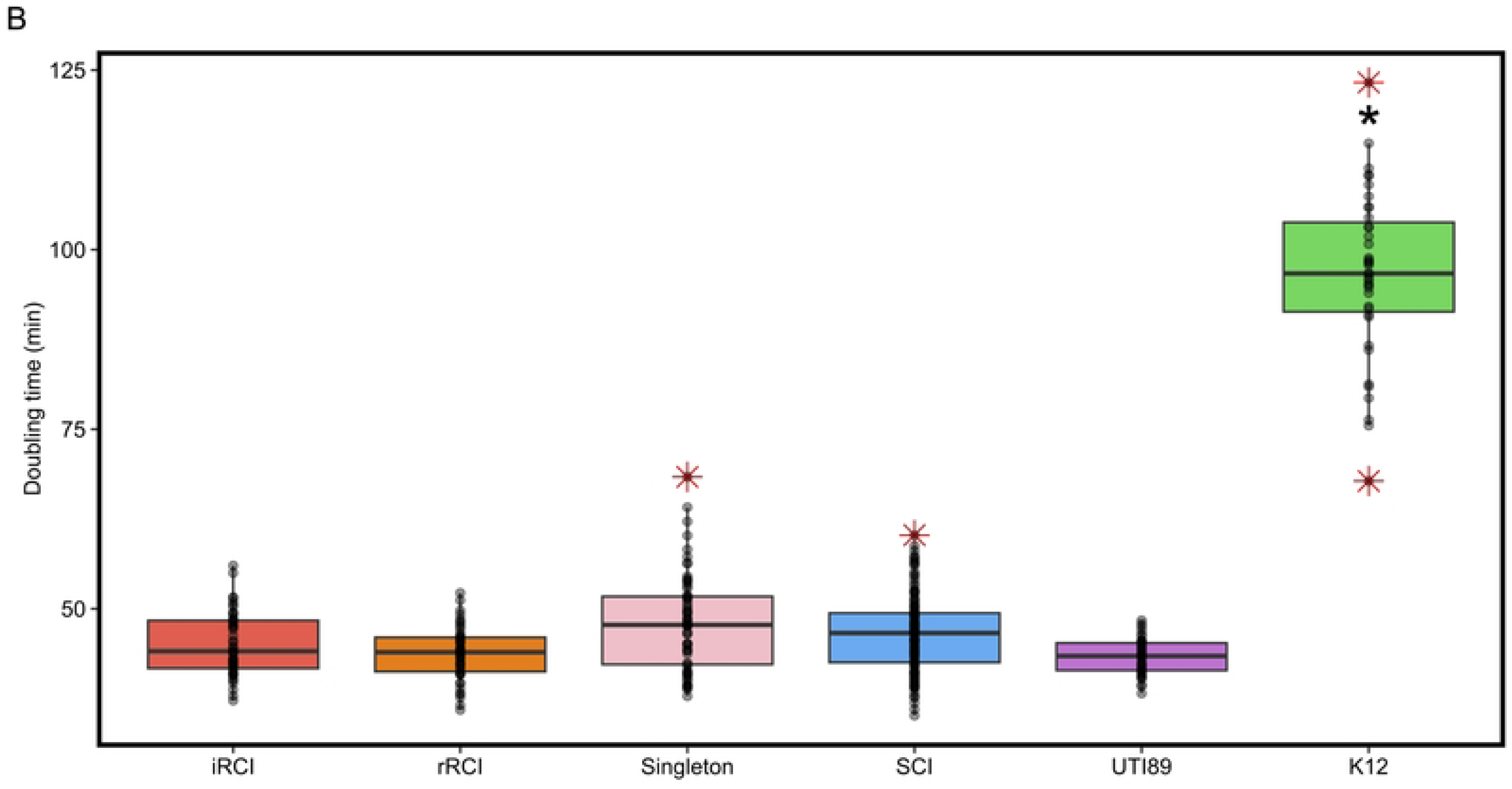
Boxplots representing the doubling time (in min) of each group of isolates in lysogenic broth (LB) (A) or in artificial urinary medium (AUM) (B). iRCIs = initial recurrent cystitis isolates, rRCI = recurrent cystitis isolate associated with relapse(s), SCIs = sporadic cystitis isolates, K12 = *E. coli str.* K12. Each dot represents a mean doubling time for one isolate in the corresponding group. Red asterisks represent extreme phenotypes for a given group (outliers).

Even though the mean doubling times in each medium were not significantly different between groups, some isolates exhibited higher or lower growth rate than the rest of their group (represented as outliers on Fig 6), with no association with specific genomic characteristics identified.

### RCIs and SCIs exhibited similar biofilm formation capacity

To evaluate whether biofilm production could be a specific trait associated with recurrence, we studied the biofilm formation capacity of 72 isolates in LB and in AUM. Globally, all groups tested exhibited low levels of biofilm formation in LB (A_590nm_ < 1) and even lower levels in AUM (A_590nm_ < 0.4). No significant differences in biofilm production were observed between groups and the positive control *E. coli* K12, neither in LB nor in AUM (Fig 7). Furthermore, no significant evolution in biofilm production was observed for RCI pairs, neither in LB nor in AUM (Fig S3).

**Fig 7.**
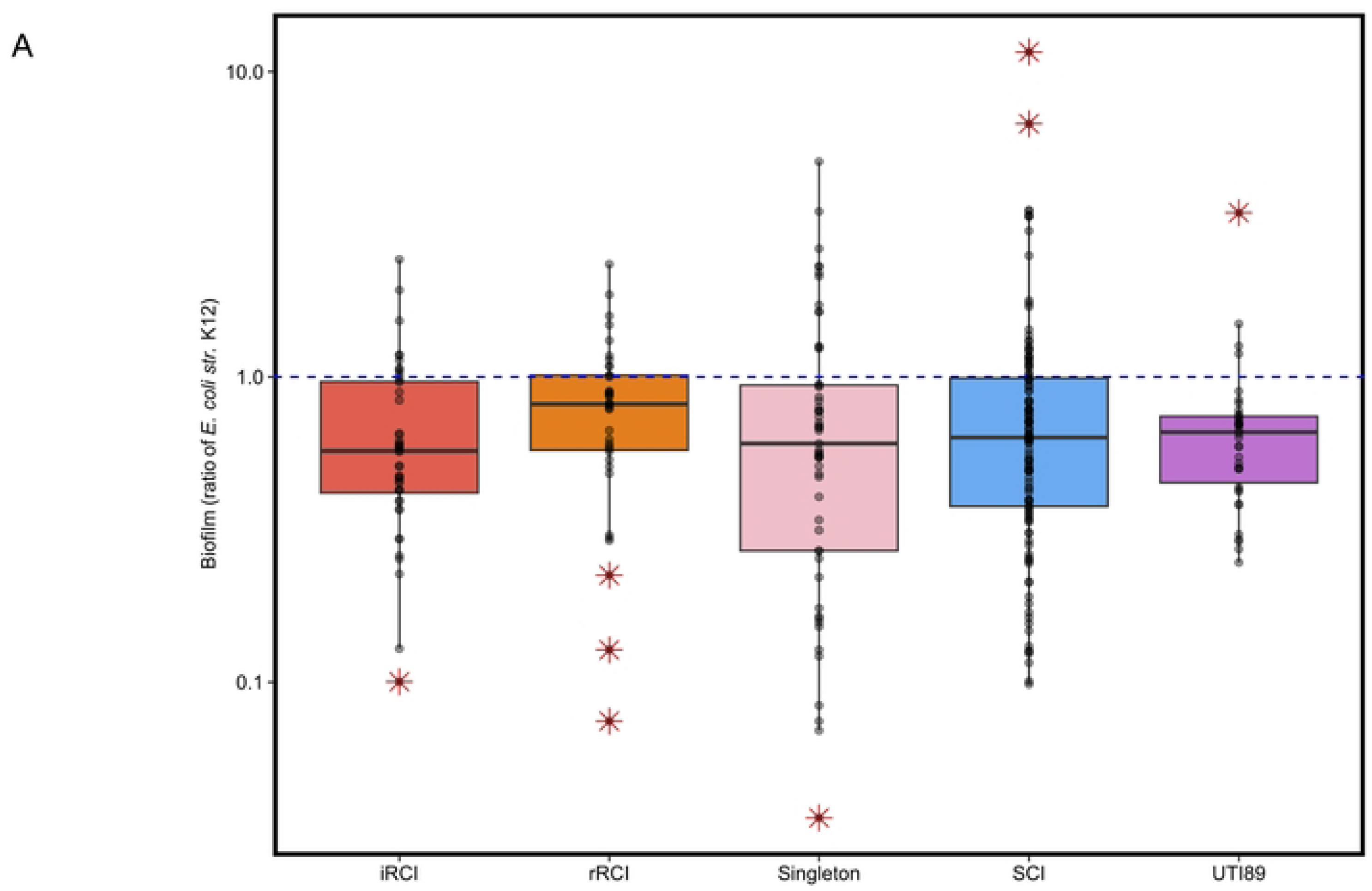
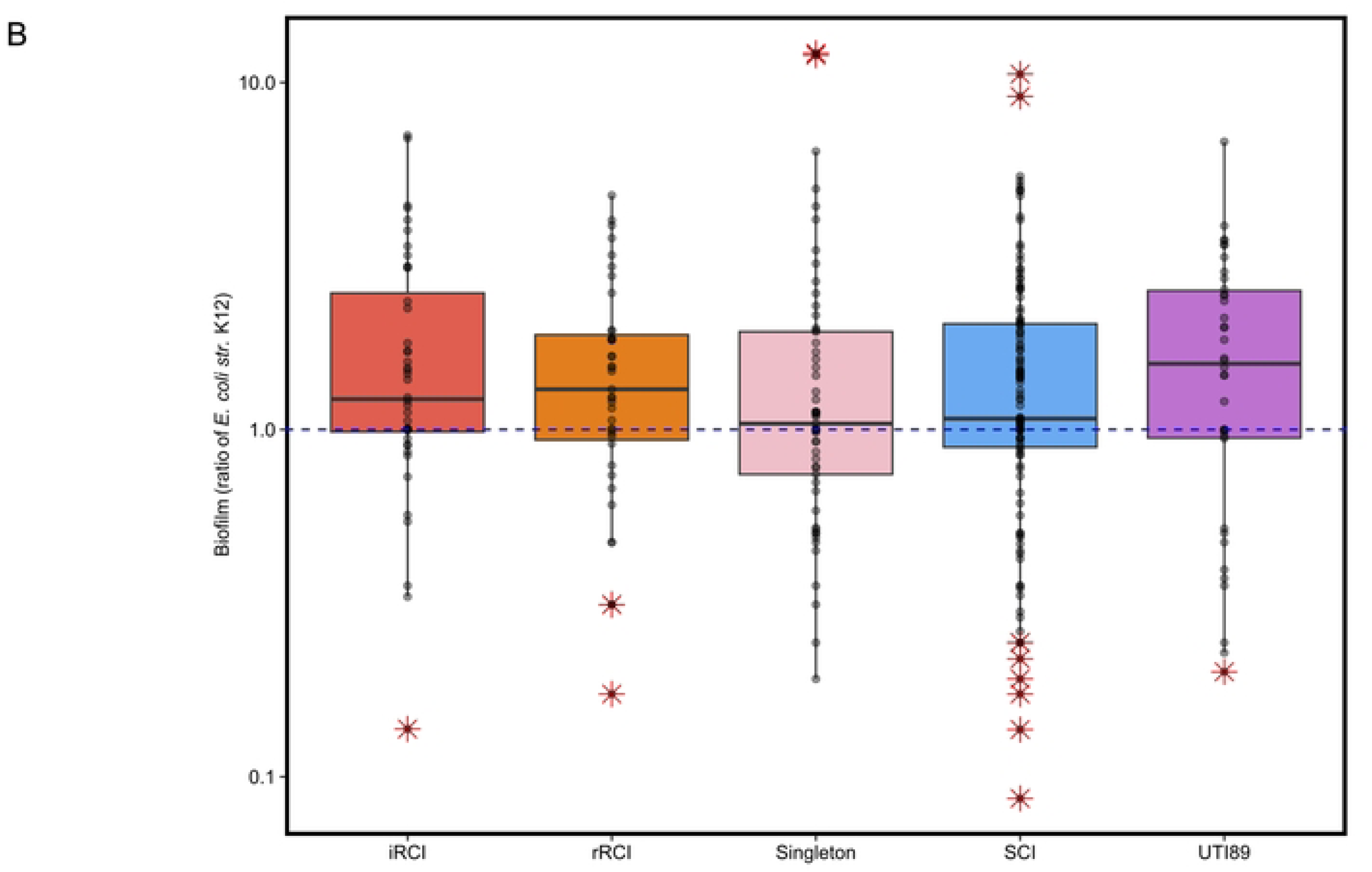
Biofilm formation in lysogenic broth (LB) (A) and in artificial urinary medium (AUM) (B) expressed as a ratio of *E. coli* str. K12 biofilm production.). iRCIs = initial recurrent cystitis isolates, rRCI = recurrent cystitis isolate associated with relapse(s), SCIs = sporadic cystitis isolates. The blue dashed line represents the biofilm production level of *E. coli str.* K12. Each dot represents a mean ratio of biofilm production for one isolate in the corresponding group. Red asterisks represent extreme phenotypes for a given group (outliers).

Of note, the overtime missense mutation identified in the PNAG biosynthesis associated poly-beta-1,6-N-acetyl-D-glucosamine synthase in rRCI_2681 (from the RCI1 series), did not lead to a significant difference in biofilm formation capacity compared to its corresponding iRCI_2630.

### Invasion of bladder epithelial cells was similar between iRCIs and SCIs

To determine whether bladder epithelial cell (BEC) invasion was a common feature among RCIs, invasion capacity of 60 isolates (10 iRCIs and their corresponding last rRCIs, 24 SCIs and 16 singletons) was evaluated by gentamicin protection assay (Fig 8). Globally, the invasion rates observed were very low (between 0.0005 and 0.1194%). Eight out of the 50 isolates (16.0%) exhibited similar or greater invasion capacity than the positive control strain UTI89 (Fig 8). The presence of intracellular bacteria was confirmed for 4 strains by fluorescence microscopy (Fig 9). There was no correlation between invasion capacity and recurrence, since the eight invasive isolates included two singleton isolates and six SCIs. Of note, the invasion rate was significantly increased between 2 iRCIs and rRCIs (RCI5b and RCI9, Fig S4). Surprisingly, *E. coli* str. K12 invasion rate was similar to that of *E. coli* UTI89.

**Fig 8.**
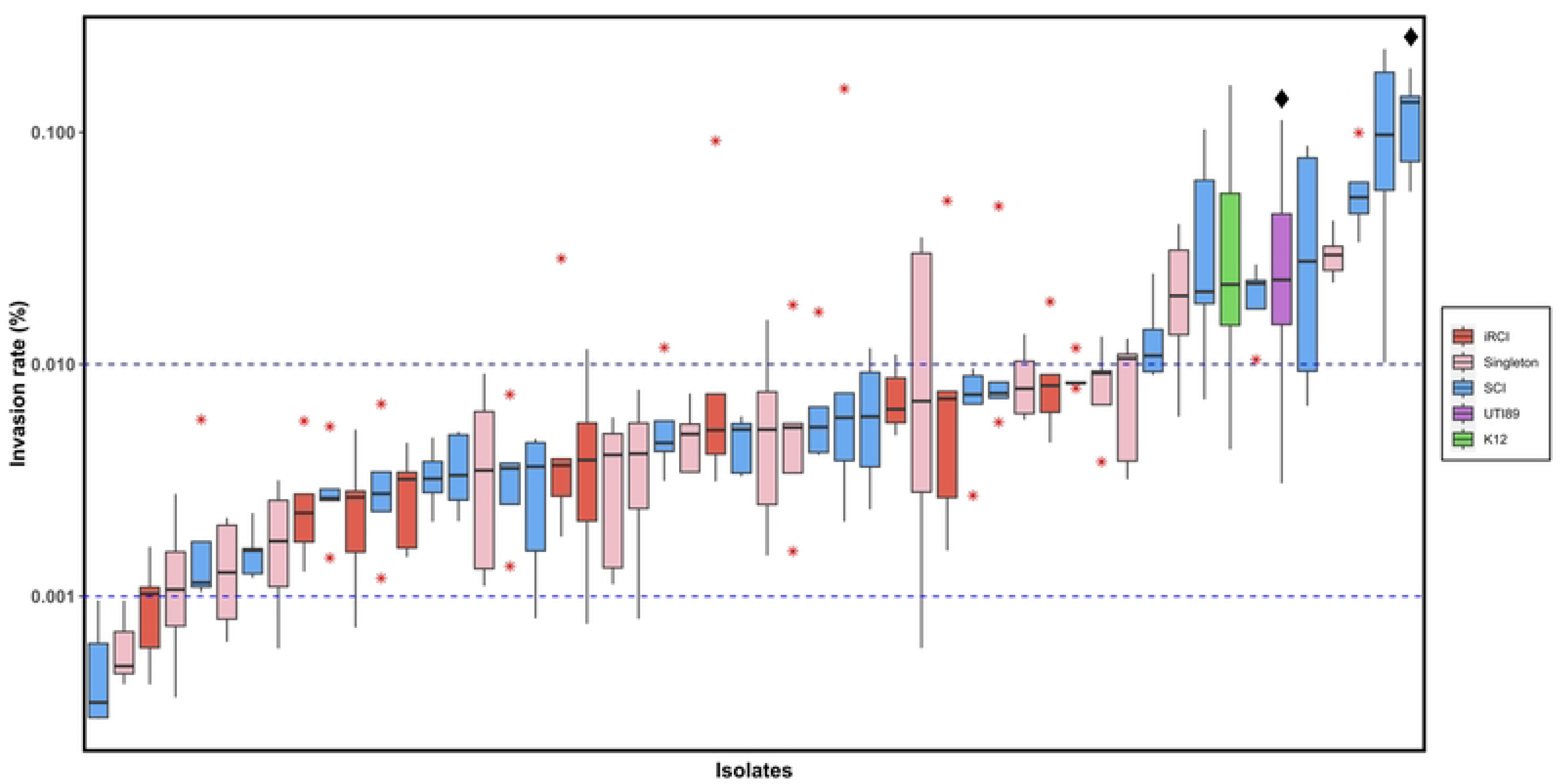
Invasion capacity of UPEC isolates in HTB-9 bladder epithelial cells after gentamicin protection assay. Boxplots represent the individual invasion rates of 10 iRCIs (initial recurrent cystitis isolates, red), 16 singletons (pink), 24 SCIs (sporadic cystitis isolates, blue), *E. coli* str. K12 (K12, green) and positive control strain UTI89 (purple). Red asterisks represent outliers. Diamonds indicates isolates for which microscopy results are presented in Fig 8. Blue dashed lines represent the invasion rate interval described by Schwartz et al. (2011).

**Fig 9.**
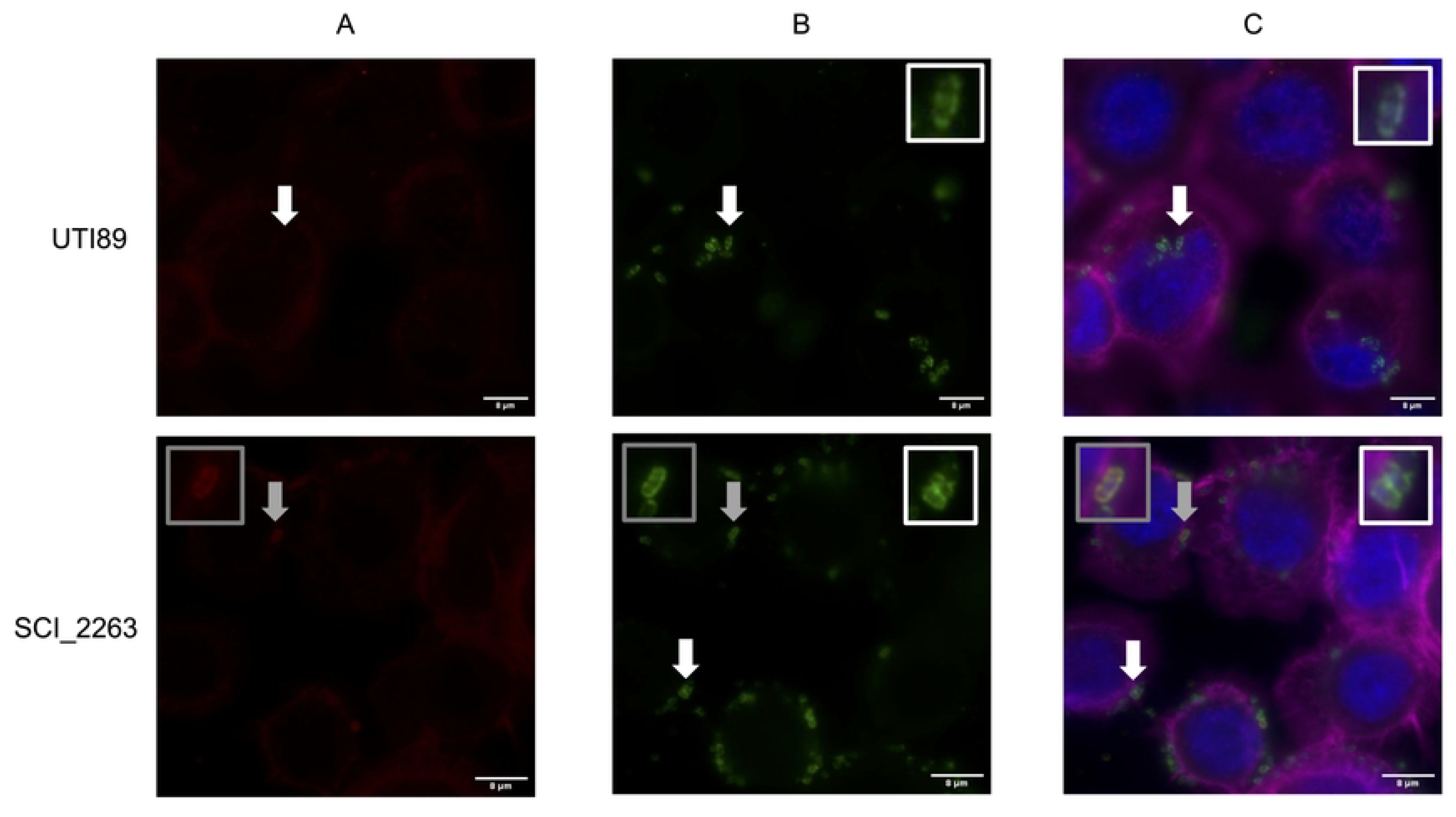
Observation of intracellular bacteria (SCI_2263, reference strain UTI89) in HTB-9 bladder epithelial cells by fluorescence microscopy. **A.** Extracellular bacteria are labelled in red (staining without cell membrane permeabilization). **B.** Total (*i.e.* intra + extracellular) bacteria are labeled in green (staining with cell permeabilization). **C.** BEC nuclei are labelled with DAPI (in blue) and actin with phalloidin (in purple). Extracellular bacteria are labelled in red or yellow (green + red) and intracellular bacteria are labelled strictly in green. White arrows and insets are focusing on strictly intracellular bacteria while grey arrows and insets on strictly extracellular bacteria.

## Discussion

Our study firstly aimed to identify *E. coli* genomic and/or phenotypic characteristics associated with UTI recurrence. To this end, we compared 24 RCIs to 24 phylogenetically paired SCIs. This pairing aimed to reduce bias due to genetic background differences and was based on CH typing, a powerful and cost-effective tool to predict MLST results and investigate UPEC [10,11].

Previous studies could not identify a specific combination of presence/absence of virulence factor genes as predictive marker of rUTI [17,18]. However, Ejrnaes *et al.* carried out a comparative study on 78 UPEC isolates causing persistence or relapse of UTI and 77 isolates followed by cure or reinfection. They reported that rUTI-associated isolates exhibit a higher aggregate virulence score, associated with 12 significantly more prevalent virulence factor genes [17]. In contrast, this study showed no difference in virulence score or VFD prevalence between groups. However, our approaches were different, Ejnraes *et al.* having worked on a larger but unpaired collection [17]. Since no difference in VFD prevalence was observed, we also studied gene polymorphism. Various levels of polymorphism were observed, but without association with recurrence. The most polymorphic genes mainly encoded adhesins and iron acquisition systems, both being already described as key virulence factors for UTI establishment [12]. This polymorphism might have a wide range of effects on bacterial phenotype, such as gene silencing, overexpression, substrate affinity modification. Such polymorphism could be the result of a host-specific adaptation. Indeed, Zdziarski *et al.* showed that the urinary tract inoculation of a single asymptomatic bacteriuria strain of *E. coli* in six patients led to genomic changes resulting in unique adaptation patterns in each patient [23].

In a recent comparative genomic analysis of 45 recurrent and 43 non-recurrent *E. coli* urinary isolates, Nielsen *et al*. did not identify any significantly associated genetic factors [18]. Unlike this study, a lower abundance of MGEs was observed in our RCI group. Of note, 8 SCIs and 2 RCIs had a prophage integrated between genes encoding an envelope stress response system involved in regulation of virulence in UPEC (*cpx* operon) and an iron efflux system (*fieF*) [24,25]. However, any effect of prophage presence was observed on growth or biofilm formation under iron limitation. The only 2 genes significantly more prevalent in SCIs derived from this prophage.

When exploring MGEs abundance, we observed that large contigs containing *tra* operon genes were present in SCIs but absent in paired iRCI, suggesting that it corresponded to plasmids, but even more interesting, plasmid curation events were also observed across intra-patient relapse series. This confirmed a result obtained by Thänert *et al.* [19], profiling the within-host adaptation of 119 lineages of UPEC sampled longitudinally from both the gastrointestinal and urinary tracts of 123 patients with UTI. Indeed, Thänert *et al.* observed that rUTI-associated isolates exhibited lower MGE richness, suggesting that MGE loss constitute a common UPEC adaptation to the urinary tract. By contrast, SCIs maintained their fitness in multiple habitats partly thanks to the conservation of their MGEs [19].

In 2021, Nielsen *et al.* showed that recurrence-associated UPECs might adapt to the urinary tract through SNP accumulation, mainly in genes involved in diverse metabolic pathways which were over-represented among the mutated genes [18]. Our results highlighted that genes affected by nsSNPs were mostly involved in various metabolic pathways (37.9%). However, this should be taken carefully since the analysis was performed on the only few mutated proteins (58 out of 160) for which a functional pathway could successfully be attributed. Moreover, metabolism genes representing approximatively 30% of the coding *E. coli* genome [26], their representation among the mutated genes of our study was not significantly higher than expected (*p* = 0.2).

Interestingly, genes involved in environmental information processing, and more specifically ABC transporters, were the second most represented functional category of genes with nsSNPs emerging in our RCI series. These ABC transporters were mainly involved in transport of important compounds for UPEC growth in urine: diverse ionic compounds (including iron, nickel and phosphate), carbohydrates and peptides [27]. Interestingly, some of the mutated genes encoding transporters were already identified as critical factors for UPEC adaptation, such as *envZ* (response to osmolarity and pH variations) [28] or *fecC* (iron transport) [29]. Such mutations combined with those affecting metabolism genes might constitute a specific adaptation to human urine composition.

No pathoadaptative genes [30] were observed in this study, probably because of the great inter-individual diversity of human urine composition [31]. Each isolate may specifically adapt its metabolism to its host, rather than all isolates broadly sharing the same adaptation to urine. This host-shaped adaptation could occur early in relapse series. Indeed, we observed that RCI evolution rate significantly decreased over the course of relapses. Moreover, overtime-conserved SNPs – which possibly represent adaptative SNPs - mostly occurred early, between the iRCI and the first rRCI.

Plasmid loss also occurred early in the course of relapses as highlighted by our data. Taken together, these results suggest that RCIs rapidly adapt to the urinary tract followed by a slower phase of micro-adaptation. However, it should be noted that the main limitation of our approach was the lack of information concerning the UTI history of the patients before inclusion. Very early adaptation events may have occurred before the iRCI sampling, leading to a possible underestimation of this early adaptation phenomenon.

One of the main characteristics of UPECs is their ability to efficiently grow in urine [27]. To our knowledge, our work is the first to assess the growth capacity of such a large isolate collection in AUM. This medium was initially described by Brooks and Keevil in 1997 and designed to reproduce the physicochemical conditions (osmolarity, pH…) encountered in urine [32] to study growth of various uropathogens including *E. coli* [33]. The main advantages of this medium compared to the pooled human urine are its reproducibility and its stability [32,34]. The higher mean doubling time observed for our clinical isolates in AUM (G = 45.8 min) than that described for UPEC in pooled human urine (between 36.3 and 40.5min) [34,35] can be explained by the greater complexity of human urine, which contains more usable substrates (carbohydrates, amino acids…) than the AUM that only contains small amounts of peptone and yeast extract [31,32].

Although no difference in mean doubling time was observed between RCIs and SCIs, we found a notable within-group diversity, that cannot be explained by the genomic profiles of the isolates. However, as genomic data did not highlight any common pattern but suggested that isolates might adapt specifically to conditions encountered in their host, the use of AUM might not be able to reveal the physiological changes following such specific adaptations. Moreover, such *in vitro* experiments do not mimic the immune pressure existing *in vivo*. Further studies should then focus on characterizing RCI growth in their host’s urine to determine whether genomic events identified in RCIs constitute a specific adaptation to the patient’s urine composition.

Biofilm has long been known to promote persistent infection in patients [36]. To date, the association of biofilm formation with recurrent UPEC UTI is controversial [17,37]. Most studies described biofilm formation in minimal media, as nutrient stress appears to be an important factor for *E. coli* biofilm induction [38]. For example, Soto *et al.* observed a positive correlation between biofilm production and recurrence in a minimal medium supplemented with LB [37]. Nevertheless, Ejrnaes *et al.* found no correlation between biofilm formation and recurrence using AB medium, a minimal medium containing 0.2% glucose and 0.5% of casamino acids [17]. In comparison to these media, AUM induces a more intense starvation. However, like in our fitness experiments, no significant differences of biofilm production were observed between RCIs and SCIs of this study, confirming the results obtained by Erjnaes *et al.* [17]. Yet, our comparison between relapsing and sporadic strains was limited by the rate of biofilm production which was very low overall. This low production of biofilm can be explained by the slower growth of isolates in AUM and could affect the quality of the results, due to the sensitivity of the detection method. As other studies have suggested that combination of mechanical and biological stresses encountered in urinary tract triggers biofilm formation [38,39], it would interesting to perform biofilm formation assays in dynamic models that better mimic urine flux.

BEC invasion by UPECs is considered as one of the main mechanisms explaining relapses [14–16]. Since its discovery, this mechanism was extensively studied in murine UTI models using the reference cystitis-causing strain UTI89 [40–42]. Data concerning clinical isolates are lacking. Using a monolayered cell infection experiments which allowed to study the formation of intracellular bacterial communities (IBCs) [43], we found that massive internalization of clinical isolates was a rare but existing phenomenon. Most RCI and SCI invasion rates were comprised between 0.001 and 0.01%, which is in the range of published data [42]. No significant correlation between relapse and invasion capacity was however supported by our data, suggesting that this factor alone cannot explain the onset of recurrence. However even though IBC is an important mechanism for UPEC persistence in bladder, it is only part of a more complex mechanism including quiescent intracellular reservoirs (QIRs) formation in transitional epithelium, which cannot be evaluated in our model. More complex models such as organoids or *in vivo* experiments should be used to further investigate the role of QIR in recurrence. Moreover, urothelial internalization is not the only mechanism that may promote UPEC persistence. The UPEC ability to durably colonize the gastrointestinal tract and vagina has also to be considered [19,44,45]. Future studies should therefore consider exploring these reservoirs through longitudinal analysis of vaginal, intestinal and bladder microbiomes of patients suffering from rUTI.

To conclude, rather than a common adaptation mechanism to the urinary tract, our results suggest a diversity of mechanisms leading to host-specific adaptation and thus to recurrence. Further studies exploring host-pathogen relationships and impact of QIR formation with organoid models in rUTI pathogenesis should be performed to contribute to translation of these results into innovative treatments.

## Material and methods

### Ethics statements

This study is based on an UPEC collection from an epidemiological study on community acquired UTI (patients prospectively included over a 17-month period), founded by the French Ministry of Health and approved by the Medical Research Ethics Committee of the Rouen University hospital (VITALE study, Clinicaltrials.gov, identifier: NCT02292160). Participating patients received an information letter and provided written informed consent.

### Bacterial isolates

This study focused on 115 UPEC isolates from the VITALE collection (Fig 1). Among them, 80 were isolated from patients with recurrent cystitis (*i.e.* with ≥ 2 episodes over a study period of 6 months or ≥ 4 episodes over 12 months). Based on a previous comparative genomic analysis, 58 isolates were involved in series of cystitis relapses (series of cystitis episodes caused by a single strain [11]) and were then defined as “relapse-causing isolates (RCIs)”. The 58 RCIs corresponded to 24 strains, each with an initial occurrence isolate (iRCIs) followed by up to 4 sequential relapse isolates per strain, for a total 34 relapse isolates (rRCIs). The remaining 22 out of the 80 isolates from patients with recurrent cystitis were responsible for a single cystitis episode during the 18-months of patient follow-up and were defined as “singletons” [11].

A control group (*n* = 35) was formed with UPEC “sporadic isolates” (SCIs) collected from patients experiencing only one cystitis episode over the 17-month period of the VITALE study. The SCIs were selected based on their genetic proximity with the RCIs, according to a phylogeny constructed with CH typing data [11]. Briefly, each iRCI was paired with a SCI from the same CH type when possible. If not, a SCI from the genetically closest CH type was used. Proximity between CH types was determined using a phylogeny based on concatenated sequences of *fumC* and *fimH* [11]. A minimum spanning tree was constructed with BioNumerics software (Applied Maths NV, Sint-Martens-Latem, Belgium) using the unweighted pair group method with arithmetic mean (UPGMA).

Comparative genomic studies were performed on 48 isolates: the 24 iRCIs and 24 phylogenetically paired SCIs from the 35 SCIs control group. Intra-host evolution studies were based on the 58 RCIs (24 iRCIs and 34 rRCIs). Phenotypic studies were performed on 77 isolates (Fig 1).

1. *E. coli* str. UTI89 and *E. coli* str. K12 were used as control strains for phenotypic experiments. *E. coli* str. K12 was purchased from the Pasteur institute (CIP 106782).

### Whole genome sequencing and assembly

Short read whole genome sequencing (WGS) and assembly were performed on SCIs and RCIs as previously described [11]. Briefly, sequencing was performed on an Illumina NextSeq500 using the Nextera XT library kit (Illumina Inc., San Diego, CA, USA) and assemblies were producted using fq2dna v21.06 (gitlab.pasteur.fr/GIPhy/fq2dna) (Table S1).

Long read WGS was also performed for the initial occurrence of each RCI (iRCI). These were amplified by overnight incubation in 10mL of lysogenic broth (LB, MP biomedicals, Santa-Ana, USA), at 37°C and under agitation. Up to 2.10^9^ cells were used for genomic DNA extraction using the Dneasy Blood and Tissue minikit (Qiagen, Hilden, Germany) according to the manufacturer recommendations for isolating Gram-negative bacteria DNA. Alternatively, the Monarch® HMW DNA Extraction kit for Tissue (New England Biolabs, Ipswich, MA, USA) was used to improve DNA fragment length. DNA concentration and quality were checked using a Qubit (Thermo Fisher Scientific, Waltham, MA, USA) and a Nanodrop 2000 (Thermo Fisher Scientific) instruments.

Up to 1µg genomic DNA was used for library preparation according to the « Native barcoding genomic DNA protocol with EXP-NBD104, EXP-NBD114 and SQK-LSK109 » (ONT, Oxford, UK). Libraries were sequenced using R9.4.1 flowcells on a minION Mk1B device. Real-time basecalling, demultiplexing and filtering were performed using MinKNOW v21.11.8 (ONT, Oxford, UK) and Guppy v5.1.13 (ONT, Oxford, UK). Quality control was then performed using Nanoplot v1.38.1 [46].

Short and long reads from Illumina and ONT sequencing, resp, were then used for hybrid assembly using Unicycler 0.4.9. Short read assemblies from the previous study [11] and newly obtained hybrid assemblies were compared using Bandage v0.8.1 [47] for visualization and BUSCO v5.2.2 [48] with the enterobacterales_odb10 lineage dataset for genome completeness evaluation. Sequencing metrics are available in Table S1.

### SCI and RCI comparative genomic study

Short read assemblies of phylogenetically 24 paired iRCIs and SCIs were annotated using Prokka 1.14.5 [49] with defaults parameters. The protein sequences of VFDs used in the study of Ejrnaes et al. [17] were retrieved from the annotated genomes and compared using MEGA X [50]. A virulence score corresponding to the total number of VFD for a given isolate. RCI and SCI short reads were also mapped to the sequence of pUTI89 (NC_007941.1) using Snippy v4.3.6 (github.com/tseemann/snippy). More than 75% coverage of the plasmid sequence was interpreted as positive match.

Pan-genome analysis of the 48 RCIs and SCIs was performed using Roary 3.12.0 [51]. Neighbor joining tree inference was performed using MEGA X [50] based on the translated core gene alignment produced by Roary. The phylogenetic tree, the core genome allele profiles obtained from the raw short reads using cgMLSTFinder v1.1.5 [52,53] and the presence/absence table produced by Roary were visualized using Phandango v1.3.0 [54].

The presence/absence table produced by Roary was also used with Scoary v1.6.16 [55] for genome wide association study (GWAS), to identify the genes that were significantly associated with recurrence. The protein sequences deriving from these genes were submitted to BlastKOALA [56] and Kegg Mapper Reconstruct [57] to classify them according to functional pathways.

The 48 annotated draft genomes were visualized using Proksee [58] and searched for phage regions between the *cpx* operon and fieF gene using Phigaro 2.3.0 [59]. Reads from each RCI were mapped to its paired SCI draft genome using snippy v4.3.6 and visualized using Proksee [58] in order to explore large sequences that were preferentially found in SCI genomes.

### Intra-patient analysis of RCI gene evolution

Hybrid genome assembly of each iRCI was annotated using Prokka v1.14.5. [49] Short reads from each rRCI were mapped to their respective iRCI annotated hybrid genome assembly using Snippy v4.3.6 and visualized using Proksee [58] to identify potential plasmid loss over time. Lost contigs were blasted (https://blast.ncbi.nlm.nih.gov/Blast.cgi) to identify whether they were previously described as plasmids, and their gene content was analyzed.

Intra-patient SNP analysis was performed by mapping short reads from each rRCI to their respective iRCI annotated draft genome assembly using Snippy v4.3.6. As a control, reads from each iRCI were also mapped to their own annotated draft genome assembly; artefact SNPs identifed this way were removed from the analysis when also found in rRCIs from the same series. The genes in which SNPs were identified between an iRCI and its rRCIs were listed and their sequences were submitted to BlastKOALA [56] and Kegg Mapper Reconstruct [57] for functional classification.

For evolution rate analysis, SNP/day rates were calculated based onthe number of SNPs obtained above for each iRCI/rRCI pairdivided by the time (days) elapsed between rRCI and iRCI sampling. Evolution rate plot was constructed using Microsoft Excel (Microsoft corporation Available from: https://office.microsoft.com/excel). Correlation between time and evolution rate was tested with Spearman’s rank correlation test, using R software (V 4.1.3, R Foundation, Vienna, Austria).

### Growth assays

The artificial urinary medium (AUM) used in the phenotypic experiments was prepared as described by Brooks and Keevil with minor modifications [32] (Table S4). Briefly, 100mM of 2-(N-morpholino)ethanesulfonic acid (MES) was added to improve buffer capacity and limit precipitate formation. Isolates were grown overnight in LB or AUM at 37°C with shaking at 150 rpm. The cultures were then adjusted to optical density (OD_600nm_) of 0.05, and 200µL of each standardized suspension was transferred in triplicate in a microtiter plate. The plate was then incubated under continuous double orbital shaking conditions (108 rpm) at 37°C during 24h in a microplate reader (Spark®, Tecan, Männedorf, Switzerland) and OD_600nm_ was measured every 15 min. All experiments were performed at least three times. The *E. coli* str. K12 substr. MG1655 (abridged *E. coli str.* K12 in the rest of the manuscript) and UTI89 strains were used as internal controls and fresh medium as negative control for each experiment. Statistical analyses of the growth curves were performed using R software (v 4.2.1, https://www.R-project.org/).

### Biofilm assays

Isolates were grown to late stationary phase in LB or AUM; overnight cultures were adjusted as described above and incubated in a microplate in static conditions at 37°C for 24h. Nonadherent bacteria were removed by washing with H_2_O before adding crystal violet 0.5%. After 10 min of incubation under gentle shaking at room temperature, the excess of dye was discarded, and each well was washed 3 times with H_2_O. Ethanol 95% was then added, and the plate was incubated under gentle shaking at room temperature for 10 min. Absorbance at 590nm (A_590nm_) was measured using a microplate reader. All experiments were performed at least three times. The *E. coli* str. K12 strain was used as a positive control and fresh medium as a negative control for each experiment. Data were expressed as percent of biofilm formation relative to the positive control.

### Cell line and growth conditions

Human bladder epithelial cell (BEC) line 5637 (ATCC HTB-9) was maintained at 37°C and 5% CO_2_ in RPMI 1640 media (ThermoFisher Scientific, Waltham, MA, USA) supplemented with 10% fetal bovine serum and 2 mM Glutamine. Cells cultures were discarded after 20 passages.

### Gentamicin protection assay

Bacterial isolates were grown in LB at 37°C for 48h in static conditions to promote type I pilus expression [43]. Bacteria were then washed three times in phosphate buffer saline (PBS) to eliminate potential secreted toxins. Bacteria were resuspended in PBS, enumerated by serial dilution on LB agar plates and used as inoculum for infections (multiplicity of infection of 15). BEC cells were seeded the day before infection in 12-well tissue culture plates at 1.6*10^5^ cells/cm^2^. Before infection, culture medium was replaced by RPMI supplemented with 5% of fetal bovine serum. To synchronize bacterial contact to host cells, plates were centrifugated at 750 rpm for 5 min at room temperature after bacterial innoculation. After 2 hours of incubation at 37°C, cells were washed twice with PBS and incubated 1 hour at 37°C in culture medium supplemented with gentamicin (50 µg/mL) to kill extracellular bacteria. Gentamicin solution was then removed and cell monolayers were washed three times with PBS before being lysed in PBS-0.2% Triton-X-100 solution for 10 min at 37°C. BEC lysates were then plated on LB agar plates and the number of CFUs obtained after a 24h-incubation at 37°C (corresponding to the number of intracellular bacteria in the initial cell lysates) was quantified. Five replicates per isolates were performed. Results were expressed as invasion percentages using the following formula:

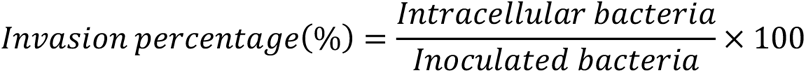

### Fluorescence microscopy

BEC cells were seeded on glass slides and infected with four UPEC isolates (UTI89, rRCI_2627, SCI_2259 and SCI_2263) as described above. After 3 hours of infection, cells were washed three times with PBS and fixed in PBS–4% paraformaldehyde. Identification of intracellular bacteria was performed thanks to a differential staining procedure [60]. Extracellular bacteria were labeled with a primary mouse anti-*E. coli* antibody (1:200; Abcam, ab35654, Cambridge, UK) in PBS–1% BSA for 1 hour at room temperature. Cells were then washed and labeled for 1 hour with a secondary goat anti-mouse antibody Alexa Fluor 546-conjugate (1:500; Invitrogen, A-11030, Waltham, MA, USA) in PBS–1% BSA. Cells were then washed with PBS and permeabilized in PBS-0.3% Triton-X-100 for 15 min at room temperature. Total bacteria were labeled with the same primary antibody and a secondary goat anti-mouse Alexa Fluor488-conjugate (1:500; Invitrogen, A-32723, Waltham, MA, USA). Cells were labeled in parallel with Hoechst 33342 (1:1,000; ThermoFisher Scientific, H3570, Waltham, MA, USA) and phalloidin conjugated to Alexa Fluor 647 (1:100; Invitrogen, A-22287, Waltham, MA, USA). Slides were mounted in Fluoromount reagent (Invitrogen, Waltham, MA, USA) and images were acquired with a Leica Thunder tissue 3D microscope and processed with ImageJ software [61].

### Statistical analyses

All statistical analyses were performed using R (V 4.1.3, R Foundation, Vienna, Austria). Comparison of proportions were performed using Pearson’s chi-squared test when applicable. Otherwise, we used Fisher’s exact test for count data. Global means comparison were performed using Kruskal-Wallis rank sum test and pairwise comparison were performed using Wilcoxon, Mann-Whitney test.

## Aknowledgements

The VITALE study (NCT02292160) was funded by the French Ministry of Health (Programme Hospitalier de Recherche Clinique) We are grateful to Normandy Region and Rouen University for funding in part the cursus of Nicolas Vautrin.

We are indebted to the volunteers. We thank all the collaborators and colleagues who helped in the study. We are grateful to the Genotoul bioinformatics platform Toulouse Occitanie (Bioinfo Genotoul, https://doi.org/10.15454/1.5572369328961167E12) for providing help and computing resources.

## Supporting informations

**Fig S1. Distribution of allelic variants of 23 virulence factor determinants in RCI and SCI groups.** Each color represents an allelic variant of the corresponding gene.

**Fig S2. Comparison of iRCI (red) and last rRCI (orange) doubling times in two media: lysogenic broth (LB) and artificial urinary medium (AUM).** Red asterisks represent outliers. Black asterisk represents significant median doubling time differences between iRCI and rRCI (p < 0.05).

**Fig S3. Comparison of iRCI (red) and last rRCI (orange) biofilm formation in two media: lysogenic broth (LB) and artificial urinary medium (AUM).** Red asterisks represent outliers. Black asterisk represents significant median biofilm formation differences between iRCI and rRCI (p < 0.05).

**Fig S4. Comparison of invasion rate of bladder epithelial cells by iRCI (red) and last rRCI (orange) in a given relapse series.** Names of relapse series are indicated above each boxplot. Red asterisks represent outliers. Black asterisks represent significant median biofilm formation differences between iRCI and rRCI (p < 0.05).

**Table S1. Sequencing, typing and pairing data for 85 isolates included in the study**

**Table S2. List and gene annotation of the 7 lost plasmids in 5 RCI series**

**Table S3. List and functional annotation of the 13 overtime conserved SNPs identified in 6 isolates from 6 RCI series**

**Table S4. Composition of AUM adapted from Brooks and Keevil (1997)**

